## Supplementary figures and images for "Fluctuations in chromatin state at regulatory loci occur spontaneously under relaxed selection and are associated with epigenetically inherited variation in *C. elegans* gene expression"

### Supplemental figure 1

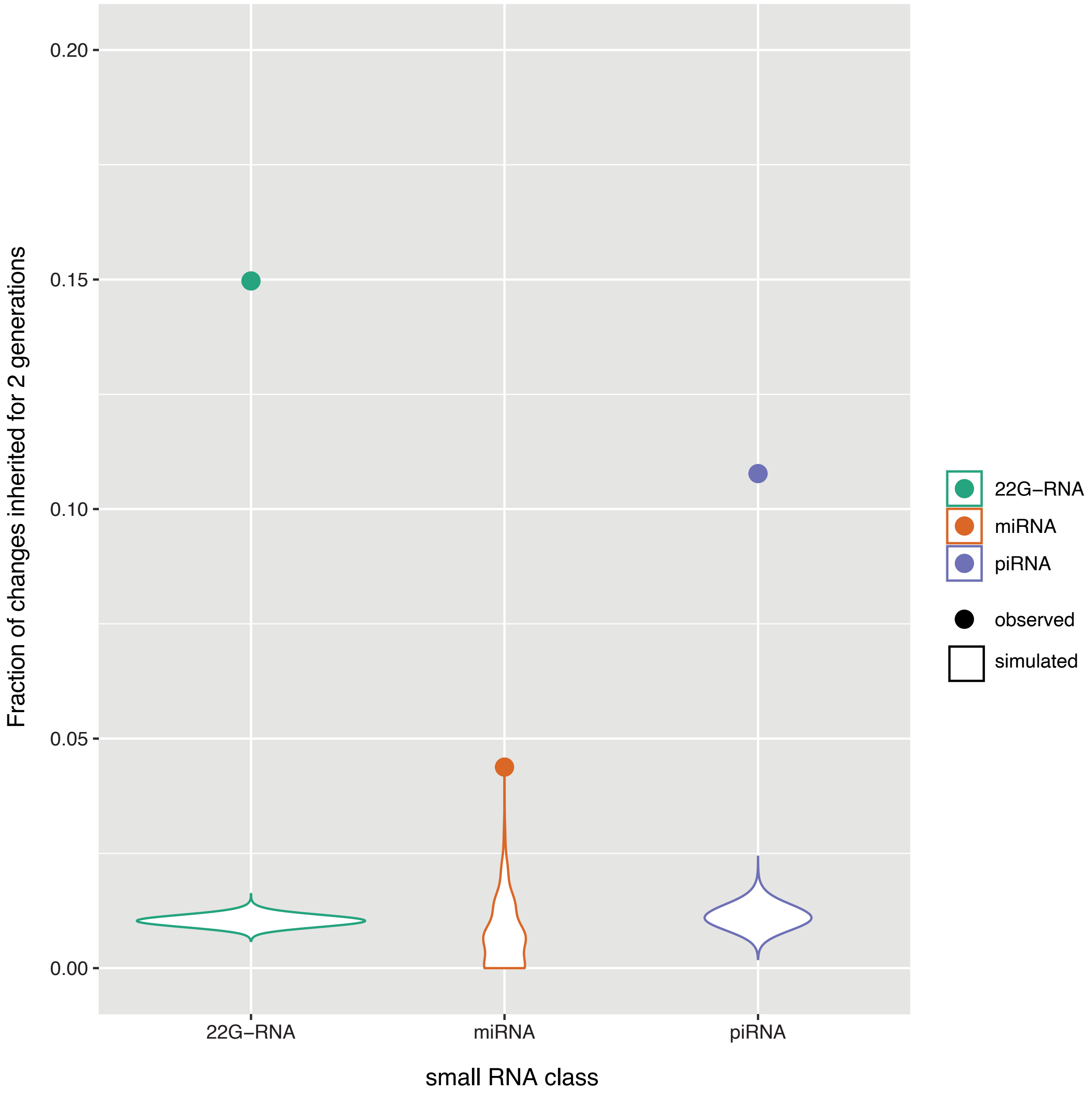


**Supplementary Figure 1**

### Supplemental figure 2

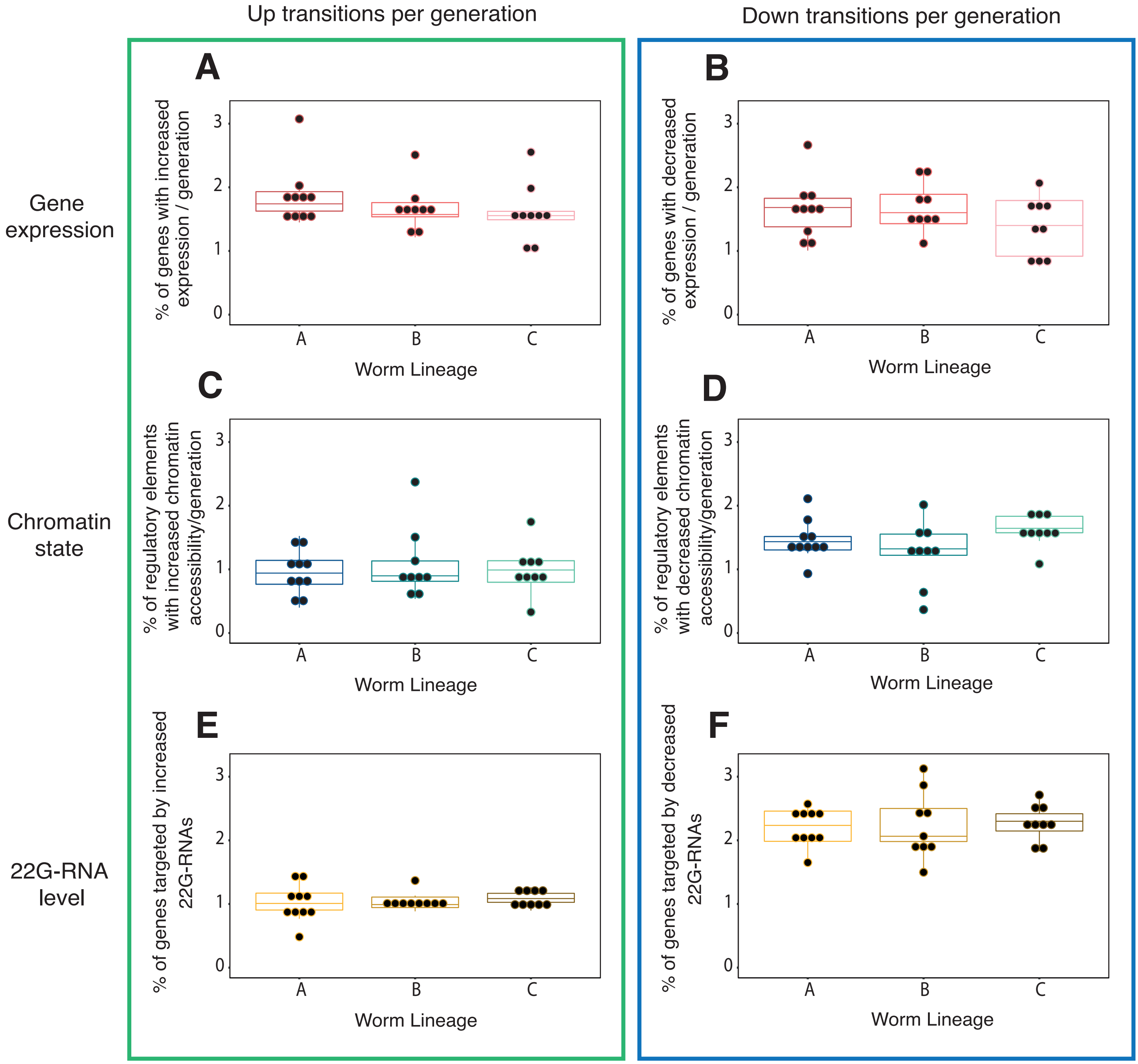


**Supplementary Figure 2**

### Supplemental figure 3

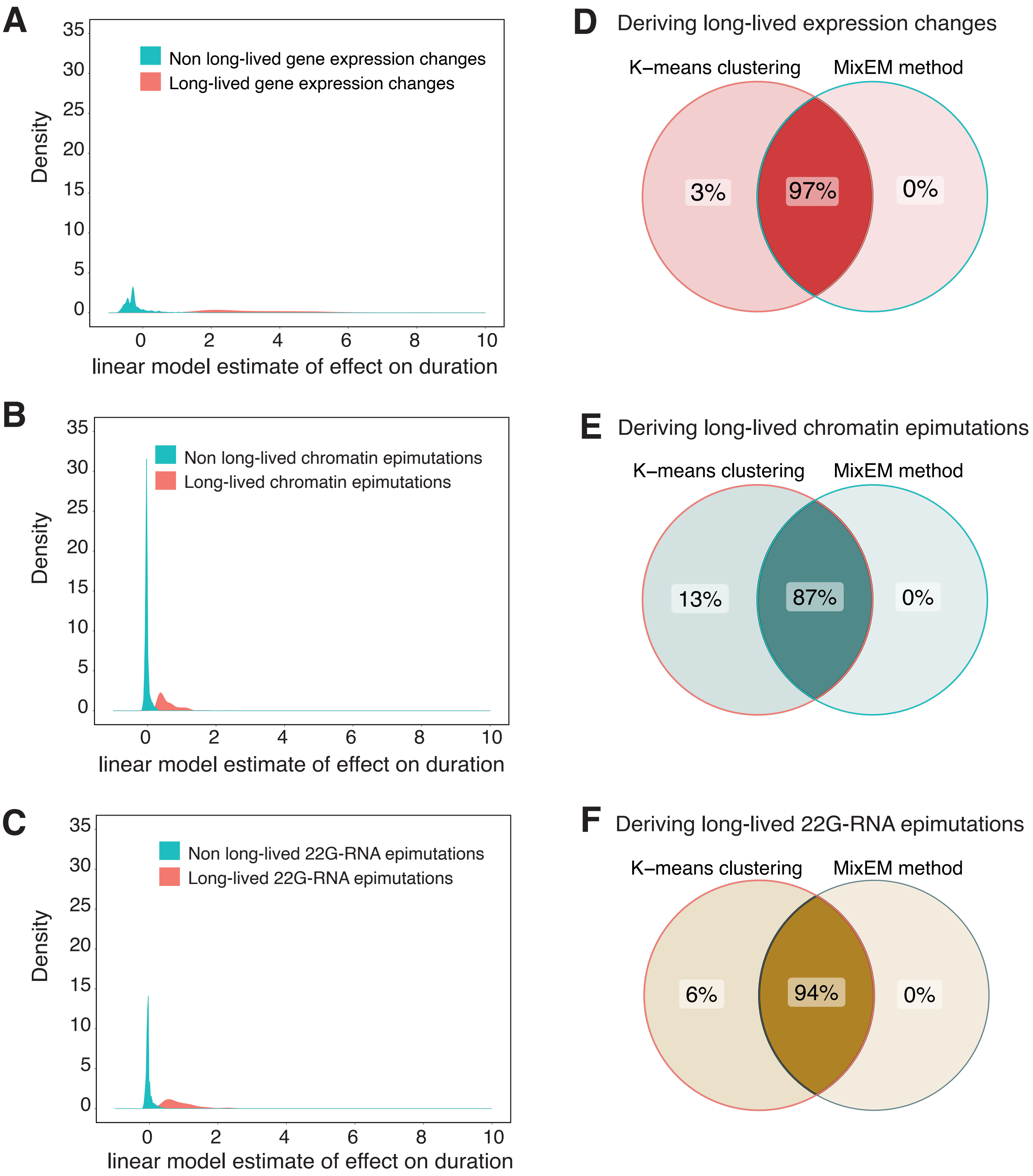


**Supplementary Figure 3**

### Supplemental Figure 4

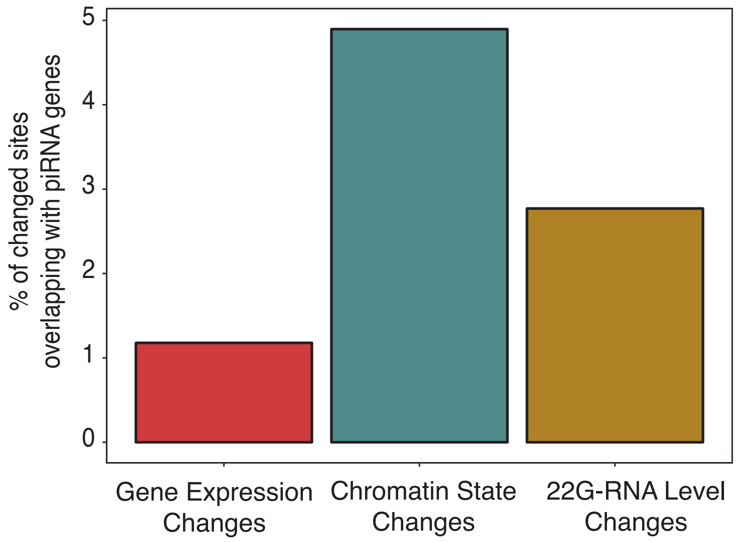


**Supplementary Figure 4**

### Supplemental Figure 5

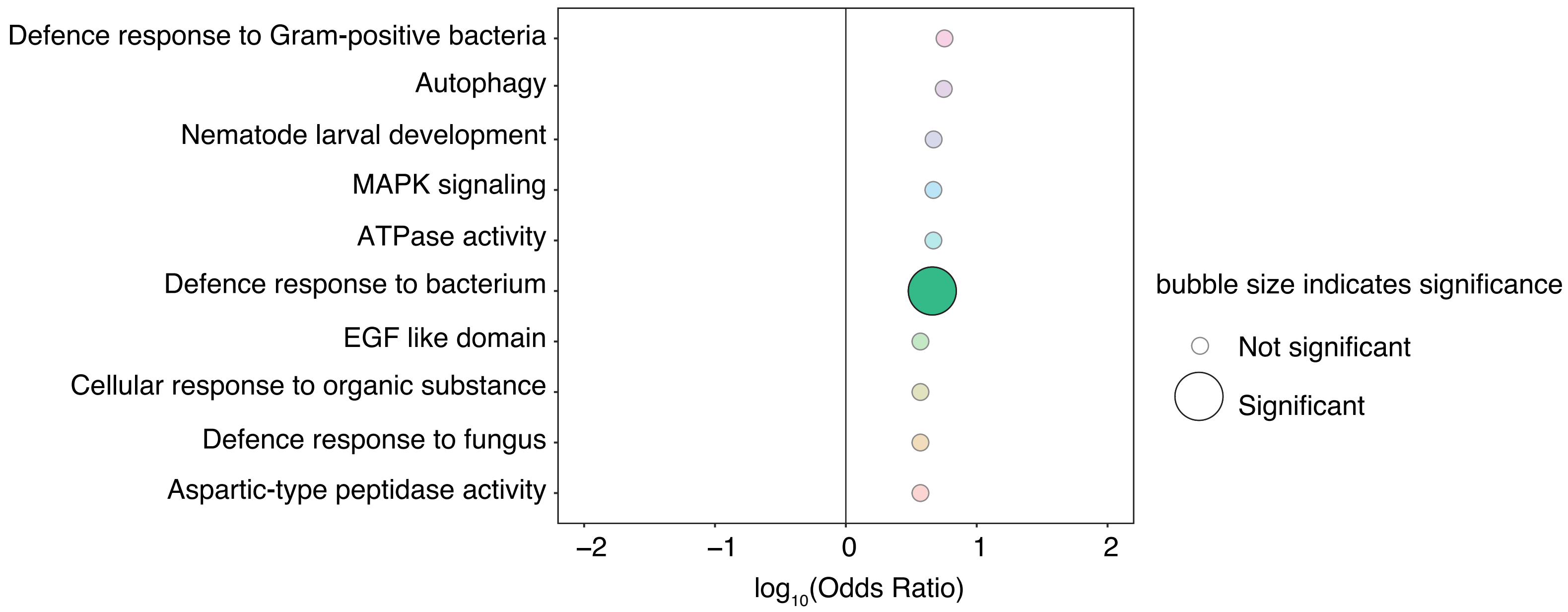


#### Supplementary Figure 5

### Supplemental figure 6

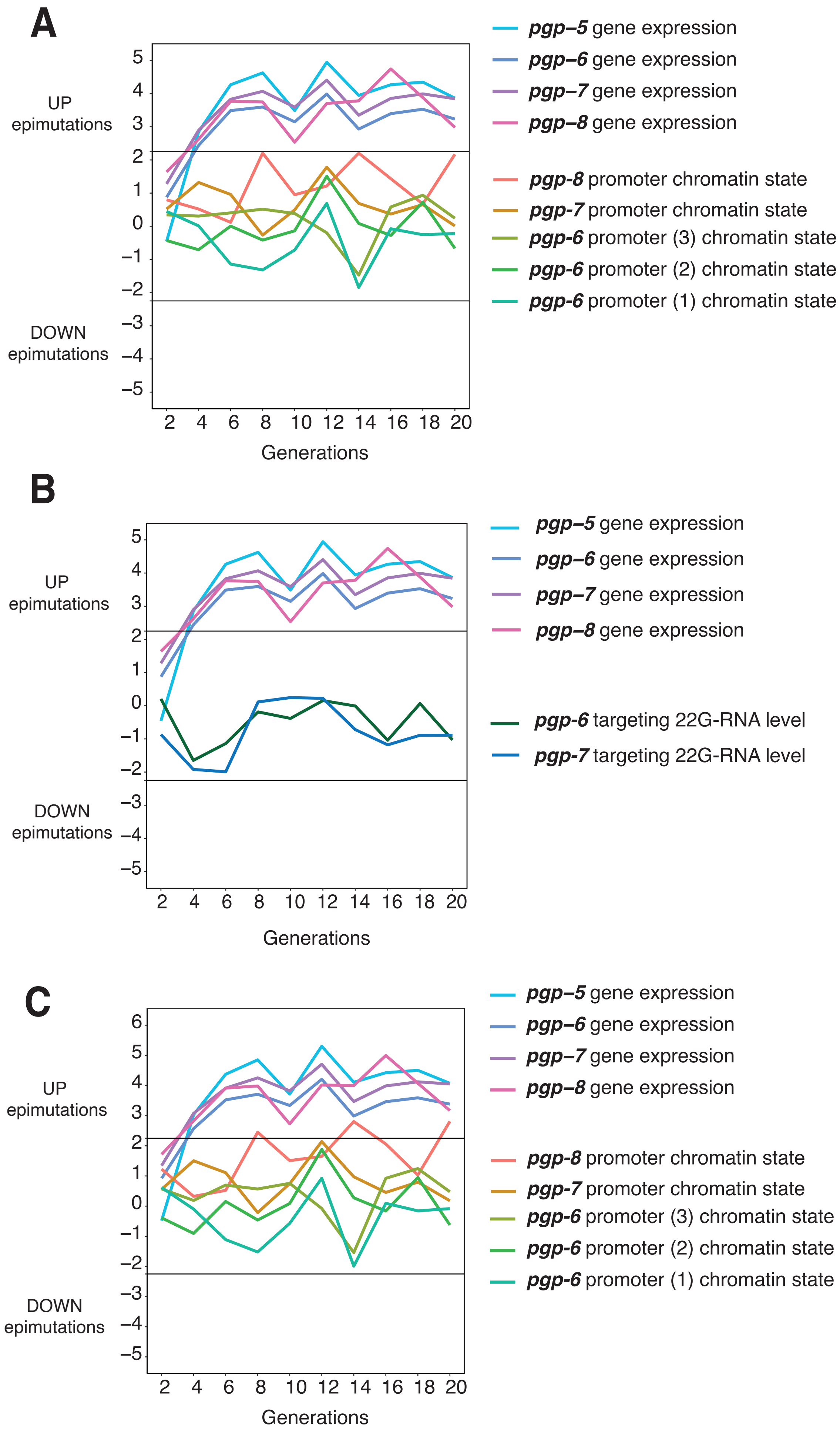


**Supplementary Figure 6**

### Supplemental Figure 7

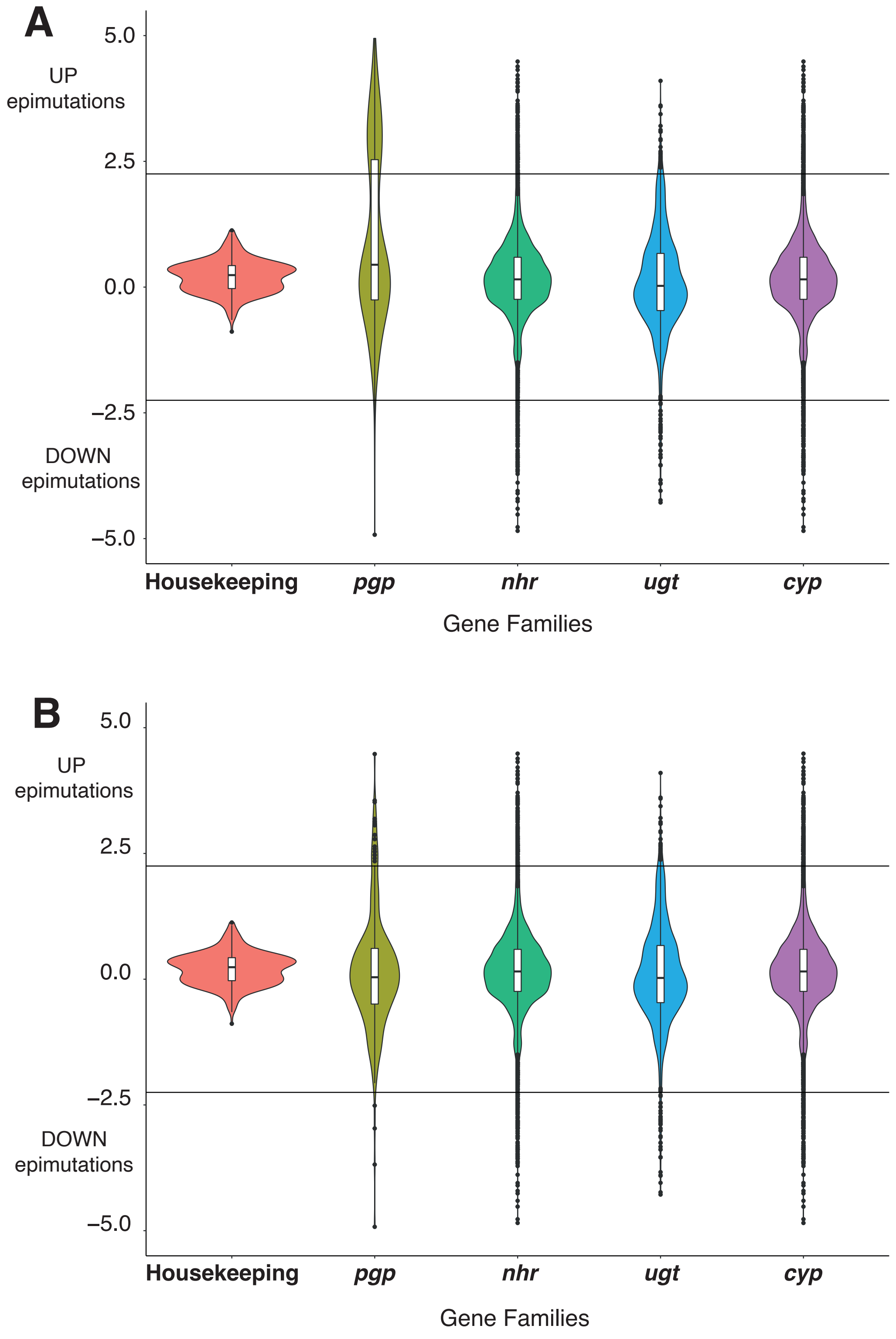


**Supplementary Figure 7**
